# Trade-off between grain yield and protein concentration is modulated by canopy photosynthesis in Japanese wheat cultivars

**DOI:** 10.1101/2022.08.17.502868

**Authors:** Keach Murakami, Hiroki Ikawa

## Abstract

A negative correlation between grain yield and protein concentration hinders an efficient nutrient supply. To analyze the causation of this inverse relationship, we compared the leaf and canopy photosynthesis of two Japanese wheat cultivars. Gas exchange measurements of leaves revealed that ‘Kitahonami’ (a high-yielding and low-protein cultivar) maintained high stomatal conductance from pre-anthesis to late grain-filling stage while stomata of ‘Yumechikara’ (a low-yielding and high-protein cultivar) were closed during the daytime, leading to decreased photosynthesis. We simulated canopy photosynthesis using a model and quantified the contribution of distinct stomatal behavior to canopy carbon gain. Although daily canopy photosynthetic gain was comparable or slightly low in ‘Yumechikara’ when model parameters obtained in the morning were used, the gain was substantially lower in ‘Yumechikara’ than in ‘Kitahonami’ when midday parameters were used. Canopy respiratory loss in ‘Yumechikara’ was greater than that in ‘Kitahonami’ during the middle of the grain-filling period because of its high canopy nitrogen content, leading to a considerable difference in net canopy carbon gain between the cultivars. Our study suggests one of the pathways for a lower carbon gain of a high protein cultivar and the greater nitrogen content does not necessarily result in a greater carbon gain.

## Introduction

Enhancing canopy photosynthetic carbon gain is believed to be a major solution for improving crop yields. Grain protein concentration is another key trait in wheat breeding programs because cultivars with high grain protein concentrations are required for bread-making. However, a trade-off between grain yield and protein concentration has been widely detected in wheat (e.g., Oury *et al*. 2003) and other cereal crops (Simmonds 1995). The inverse relationship between grain yield and protein concentration is a barrier to efficient nutrient supply systems with minimized nitrogen (N) input. Kibite & Evans (1984) concluded that the inverse relationship was not caused by genetic diversity among cultivars and suggested the involvement of N dilution by non-protein compounds. Considering that the total amount of grain N is constant, a high grain yield results in a low grain protein concentration, and *vice versa*. In their simulation based on a process model, Asseng & Milroy (2006) reproduced the inverse relationship by varying a parameter associated with the grain filling rate. It has also been suggested that competition in energy budgets for the assimilation of carbon and N shapes the inverse relationship (Munier-Jolain & Salon 2005).

Experimental data indicate that approximately 66–95 % of grain N originates from the remobilization of N accumulated in vegetative organs until anthesis (e.g., Palta & Fillery 1995; Kichey *et al*. 2007; Zhou *et al*. 2018). By monitoring radioisotope transportation in the middle of the grain-filling period in wheat, Simpson *et al*. (1983) observed that leaves, glumes, stems, and roots contributed 40 %, 23 %, 23 %, and 16 %, respectively, to the daily grain N accumulation. Martre *et al*. (2003) reported that a reduction in the grain numbers by partial ear removal manipulation significantly increased N content per grain, suggesting that grain N accumulation is source-regulated. Modern farmers produce protein-rich grains by applying high N amounts to increase aboveground biomass and N content in the grains (Zörb *et al*. 2018). In natural and agricultural systems, high N availability leads to high canopy N levels, which often exhibit a relatively high canopy photosynthetic capacity and greater biomass productivity per ground area (Chapin *et al*. 1987). Because there seems to be no distinct trend in the relationship between harvest index and grain protein concentration in wheat, we can speculate that relatively high grain yields are observed in bread wheat cultivars with substantial aboveground biomass owing to the rich N supply. Such data imply that a high canopy N content can simultaneously increase grain yield and protein concentration. What causes the inverse relationship? Triboi *et al*. (2006) affirmed that the difference in the variability of carbon and N accumulation after anthesis shapes this inverse relationship. Canopy photosynthetic carbon gain during the grain-filling period, a determinant of grain yield, varies depending on the environmental conditions during the period, whereas the total amount of grain N is largely determined by canopy N at anthesis. Although this concept explains the inverse relationship observed when canopies with the same aboveground N content are compared, it does not explain why a high canopy N content often exhibits a low yield.

A plausible reason for the observation is the fact that canopy photosynthetic carbon gain is not always proportional to the canopy N at anthesis. Simulation studies have revealed the importance of the relationship between vertical profiles of leaf N content and light environments within the canopy (e.g., Field 1983; Hirose & Werger 1987a; Anten *et al*. 1995; Hikosaka 2014). A meta-analysis revealed that the relationship between vertical profiles of N and the leaf area of the wheat canopy differs from that of other species and is closer to the theoretical optima (Hikosaka *et al*. 2016). Notably, simulation studies of canopy photosynthesis presume that leaves always perform photosynthesis according to a light response curve under controlled conditions (e.g., suitable temperature and relative humidity). The potential photosynthetic rate or photosynthetic capacity is rarely achieved under field conditions owing to various environmental constraints; therefore, the operational photosynthetic rate is usually lower than the potential rate based on the light response curve, while the respiration rate hardly decreases. Leaf photosynthetic capacity and dark respiration rate increase as a function of leaf N content (Wright *et al*. 2004). The net photosynthetic gain of a high-N canopy can be lower than that of a low-N canopy under suboptimal conditions, owing to the differences in environmental sensitivity of photosynthesis and respiration, leading to low grain yield.

Most previous studies on the inverse relationship between grain yield and protein concentration in wheat have focused on the absorption, accumulation, and remobilization of N. Although studies have evaluated the architecture and N profile of the wheat canopy (Dreccer *et al*. 2000; Bertheloot *et al*. 2008; Moreau *et al*. 2012), photosynthesis of leaf layers and canopy during grain filling has not been analyzed by associating it with the inverse relationship between grain yield and protein concentration despite its crucial roles in the grain carbon budget and. To facilitate resource-effective crop production systems, vertical profiles of leaves and N distribution in the wheat canopy should be analyzed in association with the time evolution of operational photosynthetic performance of leaf layers by comparing cultivars with contrasting canopy and grain N properties.

In this study, we analyzed the photosynthetic characteristics of the flag and lower leaves of two elite winter wheat cultivars in Japan, which exhibit a typical inverse relationship between grain yield and protein concentration. Our intensive measurements revealed that the two cultivars exhibited contrasting stomatal responses, and the low midday stomatal conductance observed in the low-yielding cultivar determined the gap between the operational photosynthetic rate and photosynthetic capacity. To quantify the effects of stomatal conductance on canopy photosynthesis, we simulated photosynthetic carbon gains of leaf layers using a simple photosynthesis model with two patterns of stomatal responses to the environment.

## Materials and Methods

### Plant material and management

Field experiments were carried out at the Hokkaido Agricultural Research Center (43.0 °N, 141.4 °E), on the northern island of Japan, Hokkaido. The site is located in Sapporo City, which faces the Sea of Japan. Because of intensive snowfall in this region, winter wheat seedlings are covered by snowpacks during winter and resume growth in mid-April.

Two winter wheat (*Triticum aestivum* L.) cultivars, ‘Kitahonami’ (KH) and ‘Yumechikara’ (YC), were used in the experiment. High-yielding KH exhibits a relatively low grain protein concentration and is suitable for the production of Japanese noodles. The soft wheat cultivar accounts for approximately 40 % of national wheat production in area basis and is the most widespread cultivar in Japan. The leaves of this cultivar are erect, which is believed to promote its high grain yield (Kasajima & Araki 2020). YC exhibits a higher grain protein concentration and lower grain yield than KH and is suitable for bread production. This extremely hard wheat cultivar accumulates a higher amount of shoot N than KH and usually forms closed canopies due to its floppy flag leaves. A typical inverse relationship between grain yield and protein concentration was observed during a four-year shading experiment conducted using the two cultivars (Shimoda & Sugikawa 2020).

Seeds were sown on 2020-09-24 using a seed drill with bulk-blending fertilizer (BB807M; Hokuren Federation of Agricultural Cooperatives, Sapporo, Japan) applied at rates of 40 kg N ha^−1^, 150 kg P ha^−1^, 85 kg K ha^−1^, and 10 kg Mg ha^−1^. KH and YC cultivars were sown at seeding rates of 180 and 250 seeds m^−2^, respectively. Eight plots were prepared for each cultivar. Topdressing was applied twice using ammonium sulfate during the vegetative stages of both cultivars. The first topdressing was applied either at the tillering stage (mid-April) or panicle formation stage (late April), and the second was applied at the flag leaf stage in late May. We broadcast 6 and 4 kg N ha^−1^ for KH, and 8 and 6 kg N ha^-1^ for YC. The doses were adopted based on the conventional protocol for commercial farmers.

Daytime and nighttime air temperatures and precipitation during the experimental period were recorded at a meteorological station adjacent to the field experiment site (Fig. 1). Heading and anthesis dates occurred 1–2 d earlier in YC than in KH. We defined heading and anthesis dates as the first days when 50% of the panicles had emerged and flowered in the field, respectively. Ear numbers at the maturity stage were 693 and 635 m^-2^ in KH and YC, respectively, which were close to the recommended values of local guidelines for the cultivars. Grain protein concentration and yield at 12.5 % (v/v) water content in KH were 8.4 % and 7.5 t ha^-1^, respectively, and 13.0 % and 6.3 t ha^-1^, respectively, in YC.

**Fig. 1.**
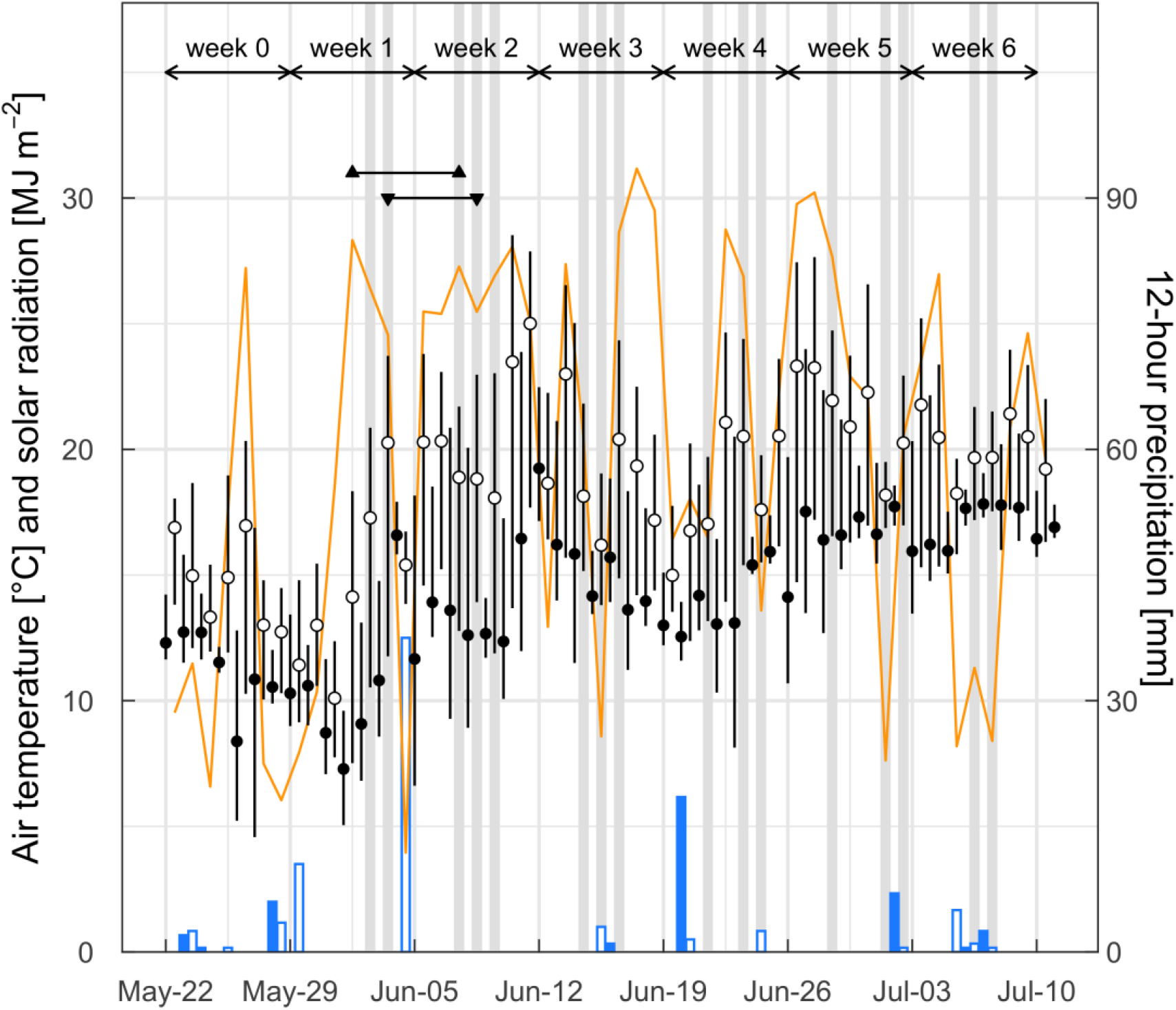
Mean, minimum, and maximum air temperatures (black), solar radiation (orange), and 12-h precipitation (blue) during the experimental period. Daytime (06:00–18:00; open symbols) and nighttime (18:00–06:00; filled symbols) temperature and precipitation values are shown. Gas exchange measurements were performed on the days indicated by gray shading. The period from heading to anthesis in ‘Yumechikara’ and ‘Kitahonami’ wheat cultivars are represented by up- and down-pointing triangles, respectively.

### Gas exchange measurements and sampling

Gas exchange measurements were performed using a portable gas exchange measurement system (LI-6400XT; LI-COR Inc., Lincoln, NE, USA). Each seedling was uprooted carefully, transplanted into a pot, and immediately subjected to approximately 1 h of gas exchange measurements in a shed. As the shed was not environmentally controlled, the air temperature and relative humidity were similar to those in the field.

We measured the light response curves of flag leaves and third leaves from the top of the plant using blue and red light-emitting diodes light sources at a photosynthetic photon flux density (PPFD) ratio of 1:9 (6400-40; LI-COR Inc.). Measurements were performed two to three times per week from early June (heading stage) to early July in 2021, when most of the leaves turned yellow. After warming up the infra-red gas analyzers, either the flag or third leaf of 8–14 plants was measured per day, as described below.

To minimize the effects of antecedent environmental conditions on photosynthetic performance, moderate actinic light was supplied for 20 min at a PPFD of 300 µmol m^-2^ s^-1^. We then increased the actinic PPFD to 1,500 and 900 µmol m^-2^ s^-1^ for flag leaves and third leaves, respectively, and recorded leaf photosynthetic parameters for 10 min. The actinic PPFD was reduced in a stepwise manner from (1500), 900, 500, 150, 75, 50, 25, and 0. Each step required approximately 5 min. Photosynthetic parameters were recorded every 10 s during the procedure, which was occasionally interrupted by ‘matching’ to calibrate the infra-red gas analyzers in charge of inflow and outflow gases. The chamber was operated at a leaf temperature of 22 °C, vapor pressure deficit of approximately 1.1 kPa, and ambient CO_2_ concentration of 400 µmol mol^-1^.

Leaves of plants subjected to gas exchange measurements were sampled to assess the leaf area and SPAD values (SPAD-502 Plus, Konica Minolta Inc., Chiyoda, Tokyo, Japan). The leaves were dried at 80 °C for three days to assess dry weight and then stored in a dehumidifier. The dried samples were cut into small pieces, and their carbon and N concentrations were measured using a CN analyzer (Sumigraph NC-Trinity, Sumika Chemical Analysis Service Ltd., Osaka, Japan). To estimate the N content of intact leaves, we obtained an equation to calculate the leaf N content per leaf area from the SPAD value (Fig. S1). Spline smoothing and estimation of the calibration line were implemented using the mgcv package in R (R ver. 3.6.2, R Core Team 2019).

### Vertical profiles of light, leaf area, and nitrogen content

The leaf area index (LAI) was estimated using a plant canopy analyzer (LAI-2200C; LI-COR Inc.) at heights of 60, 40, 20, and 0 cm above the ground on cloudy days. The incident PPFD values were simultaneously measured at the four levels using a quantum sensor (LI-190R; LI-COR Inc.) installed on the canopy analyzer. Because the heights of the bases of the flag, second, third, fourth, and lower leaves were positioned at 60–70, 40–50, 25, and < 20 cm above the ground (Fig. S2), the leaves were regarded as representative of the four leaf layers (i.e., > 60 cm, 40–60 cm, 20–40 cm, and 0–20 cm above the ground).

We calculated the light extinction coefficient (*K*_L_) and coefficient of leaf N distribution (*K*_N_) of the canopy from the vertical profiles of LAI, incident PPFD, and leaf N content, as follows:

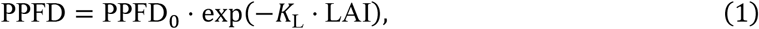

and

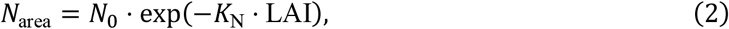

where LAI is the cumulative LAI from the canopy top to a certain level [m m^-1^], PPFD_0_ and PPFD are the PPFD values above the canopy and the level, respectively, and *N*_0_ and *N*_area_ are the leaf N content at the canopy top layer and the level [mmol m^-2^], respectively.

The vertical profiles of LAI and light were measured once every 7–9 d. The vertical profile of leaf N content was measured in samples subjected to gas exchange measurements and was estimated in intact plants based on the SPAD-N relationship (Fig. S1).

### Canopy photosynthesis model

To analyze canopy photosynthetic carbon gain, we simulated photosynthesis in leaf layers using a simple model. Based on the light response curve of a single leaf determined by the gas exchange measurement, the photosynthetic parameters of the leaf were estimated by fitting the following hyperbolic function:

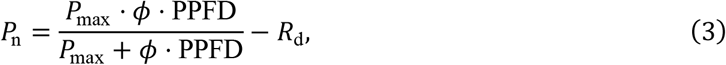

where *P*_n_ is the net photosynthetic rate as a function of the incident PPFD, *P*_max_ is the photosynthetic capacity, *ϕ* is the initial slope of the light response curve, and *R*_d_ is the dark respiration rate.

Nonlinear least-squares fitting, implemented as the nls function in R, was used to estimate the parameters.

Relationships between leaf N content and the three photosynthetic parameters were expressed as follows:

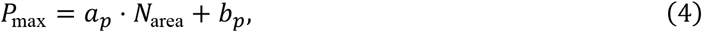

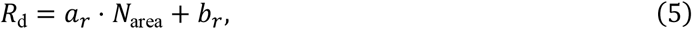

and

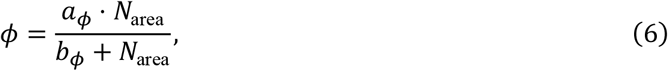

where *a* and *b* are the regression coefficients of the three parameters. For *P*_max_, four sets of regression coefficients (morning [06:00–09:00] or midday [09:00–20:00] × YC or KH) were estimated, whereas two sets of coefficients (YC or KH) were estimated for *R*_d_ and *ϕ*, respectively (Fig. S3). Robust linear fitting implemented as rlm and nonlinear least-squares fitting implemented as the nls function in R were used to estimate the parameters. The three parameters of the four leaf layers were estimated from these relationships and the leaf N profiles.

The photosynthetic rate of each leaf layer was calculated as a function of PPFD, *P*_max_, *R*_d_, and *ϕ* as shown in Eq. 3. The incident PPFD on layer *i* (PPFD_i_) was calculated as follows:

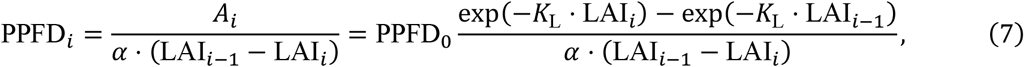

where *A*i is the flux density of photons absorbed by layer *i* [µmol m^-2^ s^-1^], *α* is the leaf light absorptance value (0.85), and LAI_i_ is the cumulative LAI from the top to the ith layer. Daily photosynthetic carbon gains of the leaf layers were calculated by integrating the products of net photosynthetic rates and LAI of the four leaf layers in a day. The canopy carbon gain was calculated by integrating these gains. To analyze the contribution of dark respiration, the net photosynthetic rates were divided into gross photosynthetic and dark respiration rates, and their integral values were calculated separately. Photosynthesis of the four leaf layers was simulated at a time step of 1 h.

## Results

### Vertical profiles of LAI, light extinction coefficient, and leaf nitrogen content

The LAI increased until anthesis around mid-June and then decreased slightly in both cultivars (Fig. 2a). LAI values were higher in YC than in KH throughout the experimental period. The light extinction coefficient (*K*_L_) in both cultivars peaked around heading, earlier than LAI, and then decreased (Fig. 2b). The *K*_L_ value was greater in YC than in KH during the grain-filling period, indicating the closed canopy architecture of the cultivar. The coefficient of leaf N distribution (*K*_N_) was close to zero in the early season because of the almost uniform greenness within the canopy (Fig. 2c). The *K*_N_ values increased until late June as the lower leaves became senescent and decreased toward zero maturity when all the leaves turned yellow. No clear differences were observed in *K*_N_ between the cultivars. The *K*_N_-to-*K*_L_ ratio showed a trend similar to that of *K*_N_, and no clear difference was observed between the cultivars (Fig. S4). The ratio was considerably lower (< 0.4) than the theoretical optimal value of 1.0 (Anten *et al*. 1995). The leaf N content of flag and third leaves, which were estimated from the SPAD-N relationship (Fig. S1), were higher in YC than in KH, particularly during the late grain-filling period (Fig. 3). Similar trends were observed for the second and fourth leaves (data not shown).

**Fig. 2.**
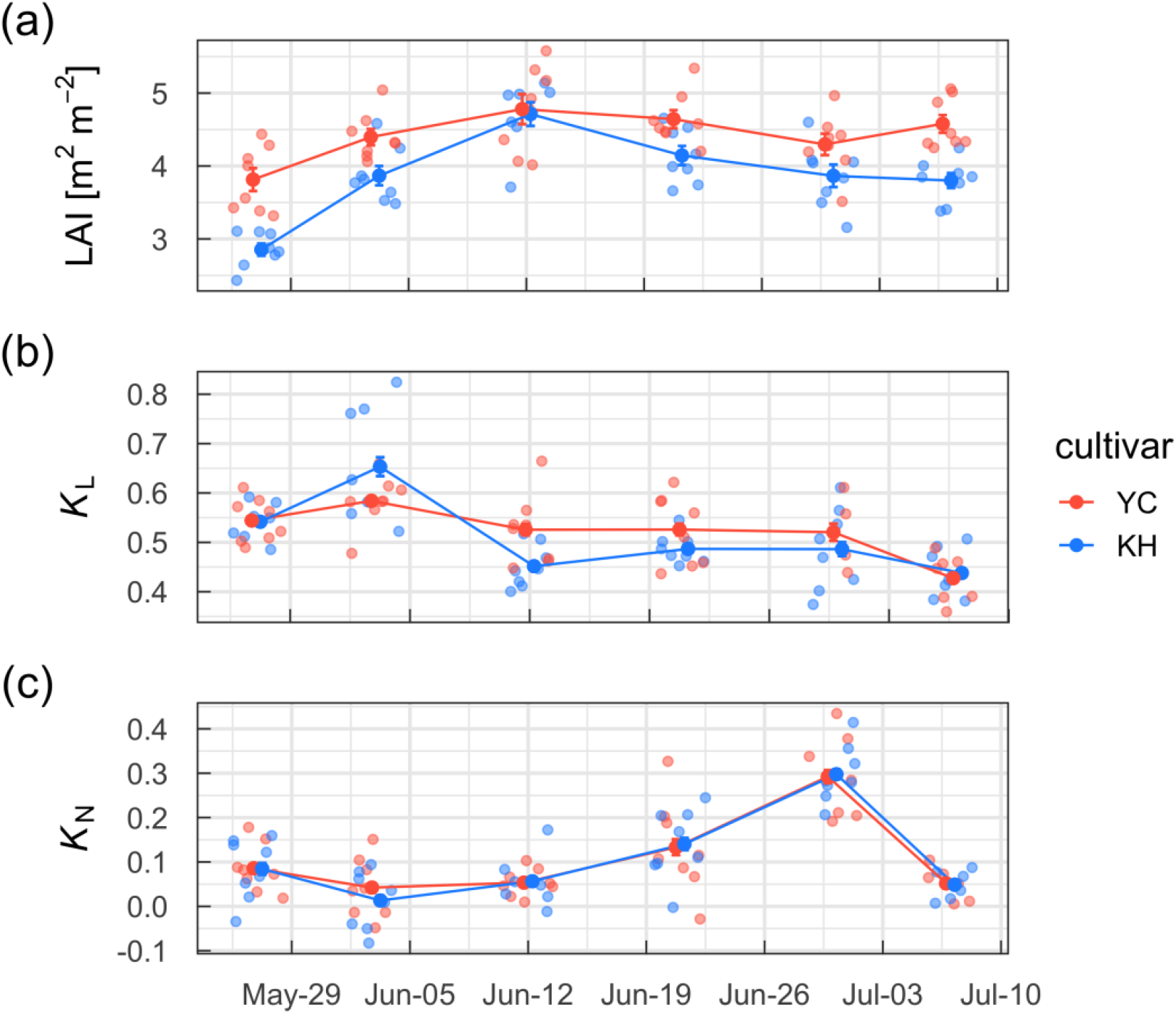
(a) Leaf area index (LAI), (b) light extinction coefficient (*K*_L_), and (c) nitrogen extinction coefficient (*K*_N_) of ‘Yumechikara’ (YC) and ‘Kitahonami’ (KH) wheat cultivars as a function of date. Values of eight replicates (small circles), their means (large circles), and standard errors (error bars) are shown.

**Fig. 3.**
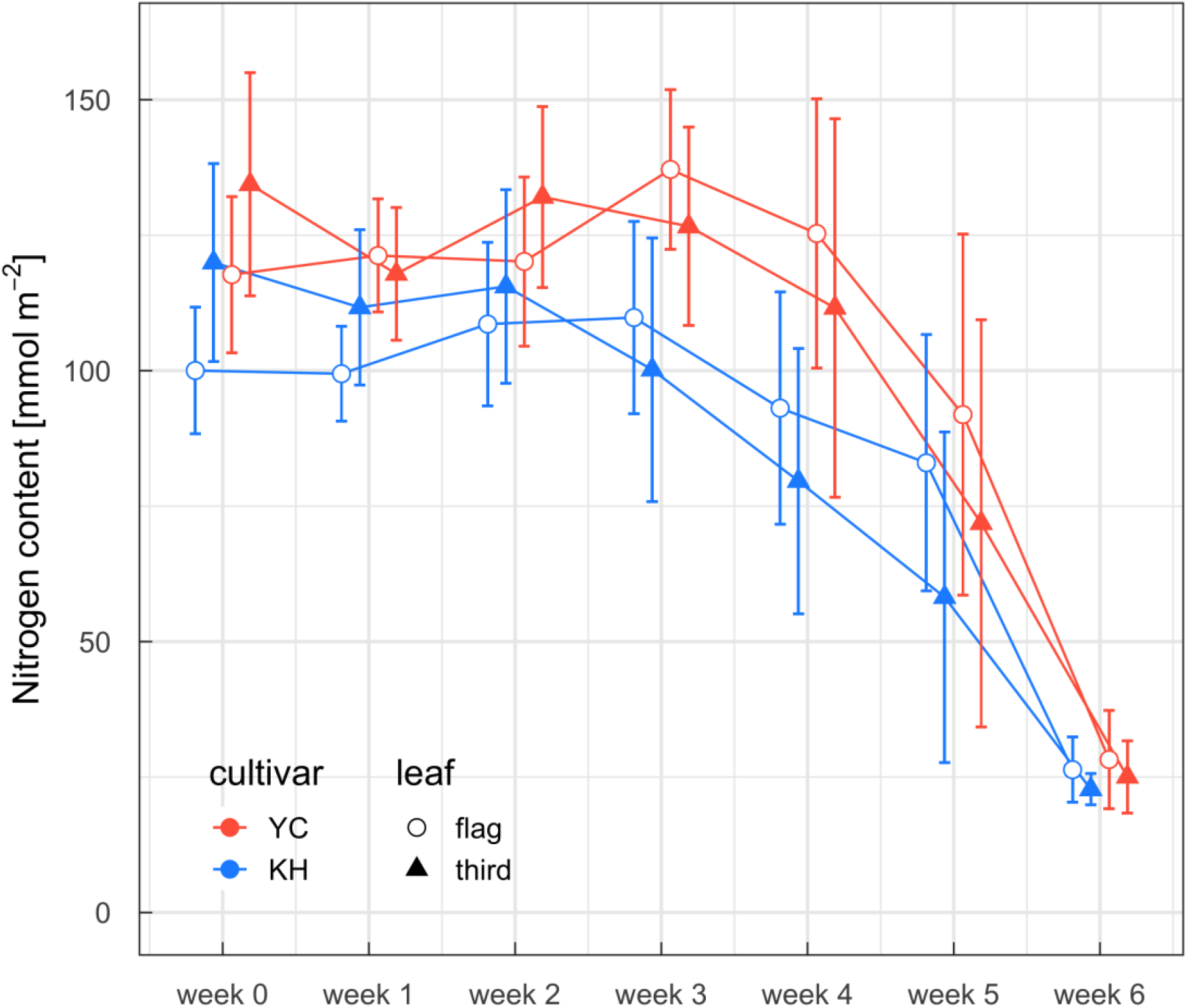
Nitrogen content of flag and third leaves of ‘Yumechikara’ (YC) and ‘Kitahonami’ (KH) wheat cultivars. Means and standard deviations are shown (*N* = 32–133). Values were estimated from a relationship between SPAD values and nitrogen content (Fig. S2).

### Photosynthetic performance of flag and third leaves

The net photosynthetic rate of KH leaves tended to be higher than that of YC leaves irrespective of PPFD during the initial growth period (Fig. 4a). A similar trend was observed for the flag leaves until the leaves became progressively senescent. In the third leaves, the difference was not obvious during the later growth period, which was due to the rapid decrease in N content of KH (Fig. 3). We also confirmed that the net photosynthetic rate of flag leaves was lower in YC than in KH based on the gas exchange measurements of intact leaves in the field under dry conditions on 2022-06-14 in the subsequent season (Fig. S6). The difference in the net photosynthetic rates seemed to be caused by the higher stomatal conductance in KH than in YC (Fig. 4b).

**Fig. 4.**
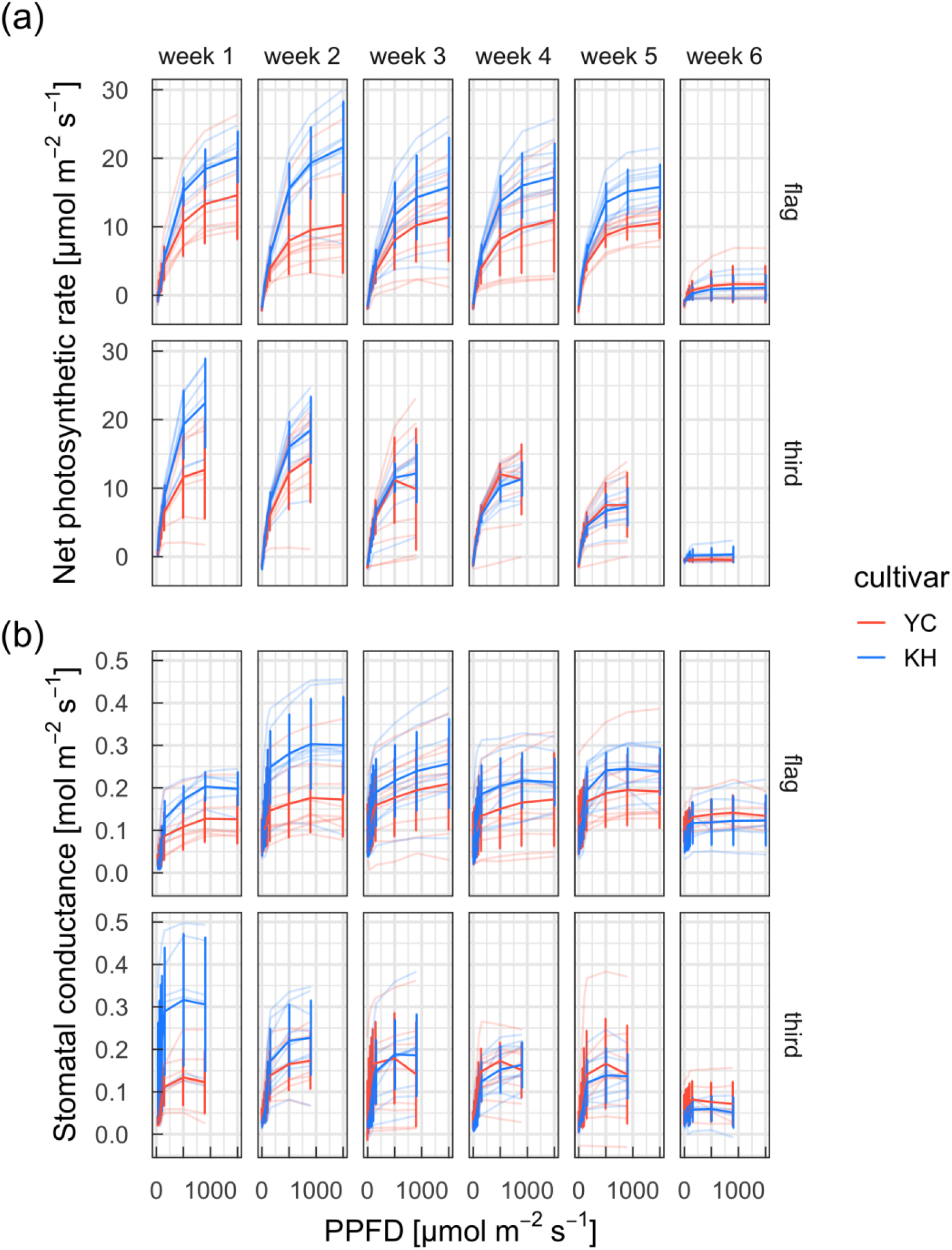
Net photosynthetic rates and stomatal conductance of flag and third leaves of ‘Yumechikara’ (YC) and ‘Kitahonami’ (KH) wheat cultivars as a function of photosynthetic photon flux density (PPFD) in different weeks. Values of samples (thin lines), their means (thick lines), and standard deviations (error bars) are shown (*N* = 5–11).

We noticed that YC exhibited a low photosynthetic rate and stomatal conductance during the daytime. To assess the effects of time of day on photosynthetic performance, we plotted light response curves from data obtained early in the morning (06:00–09:00) and at midday (11:00–14:00) (Fig. S5). A slight difference was observed in the effect of time on photosynthetic performance between the cultivars in the morning, suggesting that variations in their biochemical characteristics were minor. However, the photosynthetic rates of YC were lower than those of KH at midday. The diurnal course of stomatal conductance indicated that KH stomata remained open at midday while YC stomata were closed (Fig. 5).

**Fig. 5.**
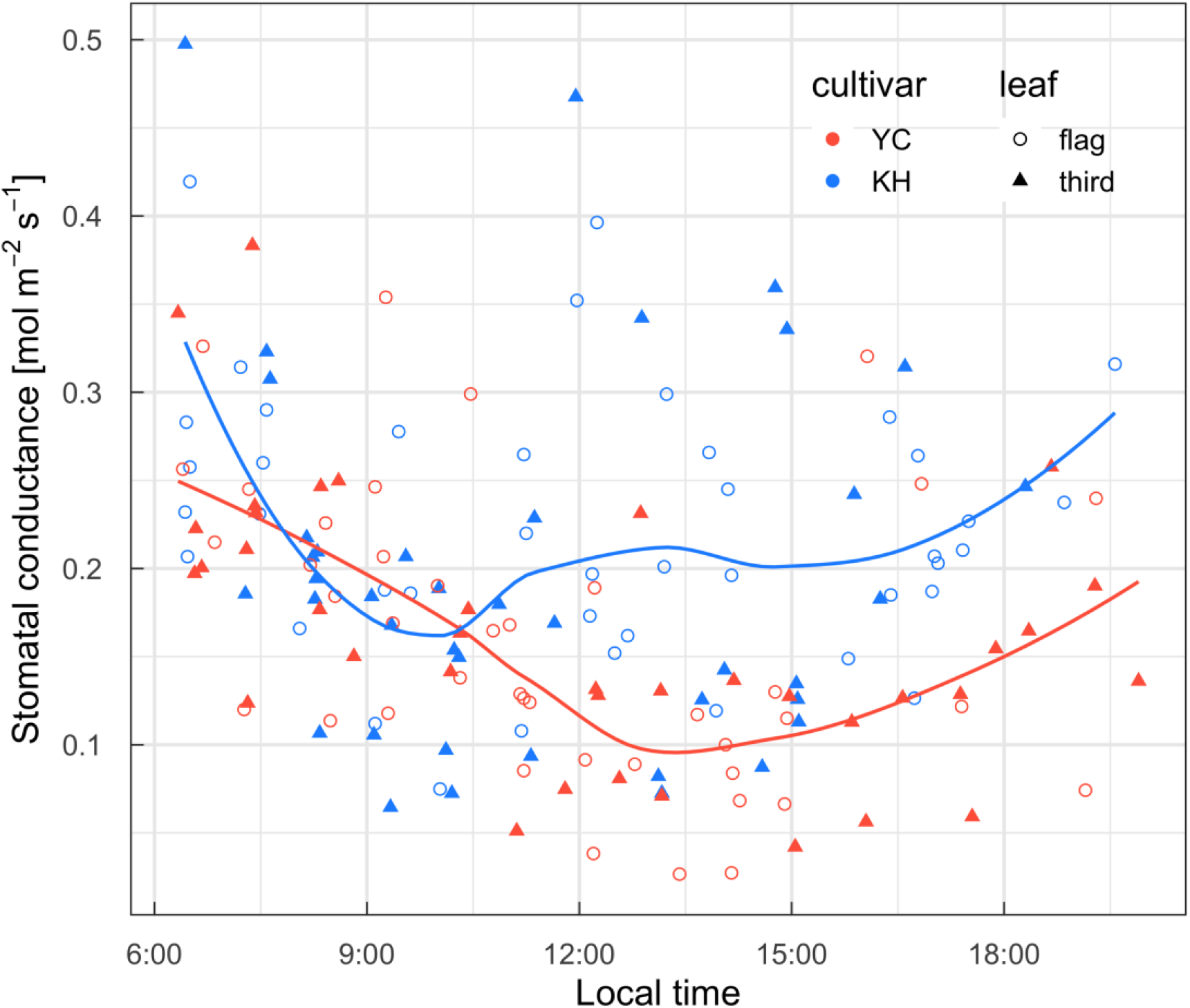
Diurnal course of stomatal conductance of flag and third leaves of ‘Yumechikara’ (YC) and ‘Kitahonami’ (KH) wheat cultivars. Values obtained at a photosynthetic photon flux density of 500 µmol m^−2^ s^−1^ are shown. Values obtained over the last week (week 6) were not used because of progressive leaf senescence. Regression lines are smooth splines.

### Simulated canopy and leaf layer photosynthesis

To assess the effects of the distinct photosynthetic light responses in the morning and at midday on the carbon budget during the experimental period, we simulated daily canopy photosynthetic carbon gains using a simple canopy photosynthesis model consisting of four leaf layers. Whole canopy photosynthetic carbon gain fluctuated significantly depending on meteorological conditions, particularly radiation (data not shown), until late June, and then dropped to approximately zero in July due to progressive leaf senescence (Fig. 6). The gross daily carbon gains simulated using the model parameters obtained in the morning (i.e., morning-type photosynthesis) were lower in YC than in KH except for a few days in late June (Jun-20 to Jun-26). A significant difference was observed in carbon gains between the cultivars particularly in the midday-type photosynthesis, which could be attributed to the contrasting stomatal behaviors of the cultivars. Whole canopy respiration loss peaked when the canopy LAI and leaf N content reached their maximum values (Figs. 2a and 3). Respiration loss was greater in KH at its peak and greater in YC during the late grain-filling period. The net photosynthetic carbon gain of the canopies (i.e., gross carbon gain − respiration loss) was always lower in YC than in KH because of the relatively high respiration rate during the late grain-filling period. The net carbon budget of YC over the last week was negative because of the rapid reduction in gross carbon gain and hardly affected respiration loss. The results indicate that the photosynthetic radiation use efficiency was lower in YC than in KH despite its high LAI.

**Fig. 6.**
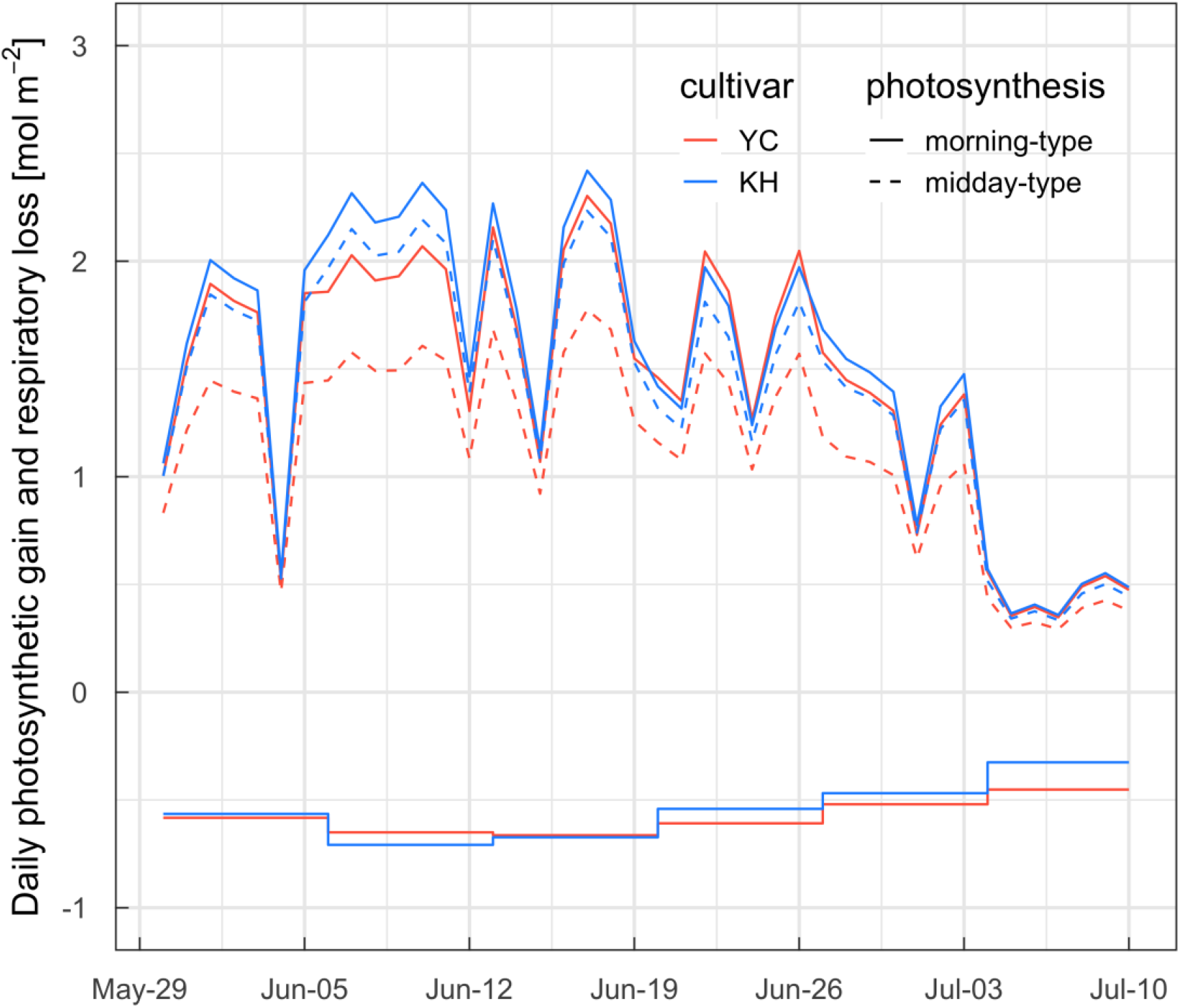
Daily simulation of whole canopy gross photosynthetic carbon gain (upper lines) and respiratory loss (lower stepped lines) in ‘Yumechikara’ (YC) and ‘Kitahonami’ (KH) wheat cultivars. Estimates presuming that leaves exhibited morning- or midday-type photosynthetic light responses throughout the day are shown.

To further analyze canopy photosynthesis, cumulative photosynthetic carbon gains of leaf layers in both cultivars and photosynthetic types were simulated (Fig. 7). Provided that leaves maintained the morning-type photosynthetic light response throughout the experimental period, the cumulative carbon gain of YC flag leaves was higher than that of KH leaves, which could be because of the high N content in the upper layer of YC canopy (Fig. 3) and the greater interception of incident light of this layer (i.e., high *K*_L_; Fig. 2b). In contrast, carbon gain was higher in KH than in YC at all leaf levels in the midday-type photosynthesis, which was attributable to the smaller stomatal conductance and high respiration rates in YC. The fourth leaves of YC served as a carbon sink rather than a carbon source when the leaves maintained the midday-type photosynthesis.

**Fig. 7.**
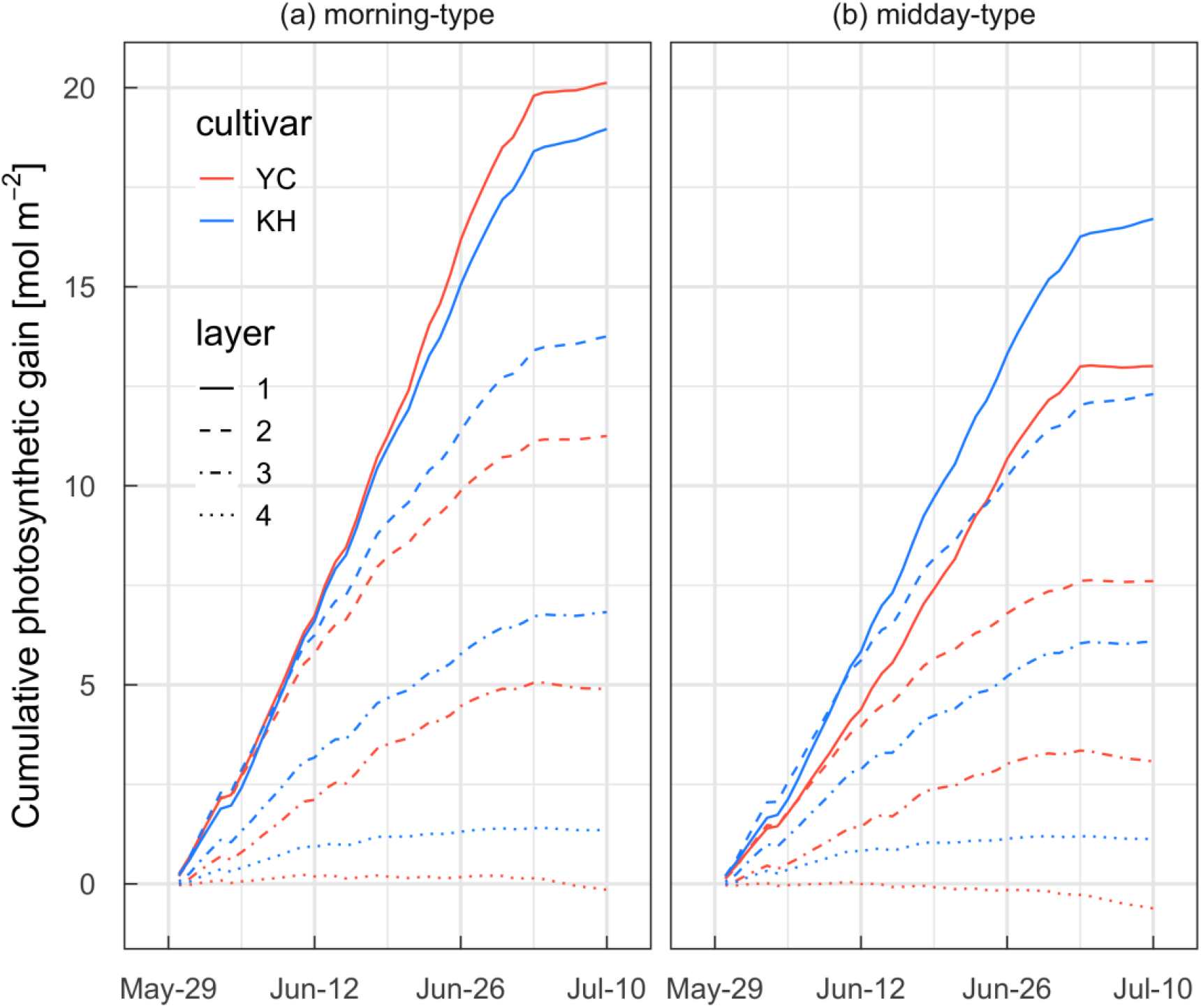
Simulation of cumulative photosynthetic carbon gain of leaf layers of ‘Yumechikara’ (YC) and ‘Kitahonami’ (KH) wheat cultivars. Estimates presuming that leaves exhibited morning- or midday-type photosynthetic light responses throughout the day are shown.

## Discussion

### Photosynthetic carbon gain of high canopy N is influenced by environmental factors

Because N is one of the most limiting resources for plant growth in both natural and agricultural systems (Chapin *et al*. 1987), high N availability often results in high biomass production per ground area. High canopy N also promotes protein accumulation in grains, leading to high grain protein concentration. Studies comparing extreme N regimes have showed that high-N treatments result in higher yield and protein concentration in wheat grains than low-N treatments (e.g., Triboi *et al*. 2006; Barraclough *et al*. 2014). However, a comparison of canopies with substantial N application among genotypes, years, sites, and/or environments has shown that high grain protein concentrations originating from high canopy N accumulation often result in low grain yields. Although previous studies have investigated the mechanisms underlying the inverse relationship between grain yield and protein concentration, with a focus on absorption, accumulation, and remobilization of N (see also the Introduction), the involvement of canopy photosynthesis in this relationship has not been comprehensively analyzed. The present study revisited this relationship by analyzing leaf- and canopy-level photosynthesis and observed that canopy photosynthetic carbon gain of the high canopy N was lower than that of the low canopy N, which partially explains the inverse relationship. The low canopy carbon gain in YC, a low-yielding and high-grain protein cultivar, was attributed to 1) a low net photosynthetic rate due to stomatal closure during the daytime and 2) a high respiration rate of the high-N canopy. We are not certain whether a low net photosynthetic rate due to stomatal closure during the daytime is unique to the YC cultivar or whether it is a common trait of hard-wheat cultivars. No study has investigated the correlation between the two factors. A high respiration rate may apply to other high-grain protein cultivars because they often require more N application to ensure high grain protein concentrations. In the present study, we did not observe notable differences in *K*_N_ and *K*_N_/*K*_L_ ratio between the cultivars (Figs. 2b and S4). The results suggest that variations in the yields of the cultivars could not be attributed to the inconsistencies in the vertical profiles of leaf N distribution and light.

In general, the high leaf N content is associated with high photosynthetic capacity (Evans 1989; Wright *et al*. 2004). The gross photosynthetic rate does not correspond to its potential capacity when any of the factors are not optimal because it is limited by various environmental factors (e.g., PPFD, air and leaf temperatures, and vapor pressure deficit). In contrast, dark respiration rate mainly depends on leaf N content (Hirose & Werger 1987b; Makino & Osmond 1991; Wright *et al*. 2004) and is less affected by short-term environmental conditions, except for temperature response. Under suboptimal conditions for photosynthesis, the performance of leaves with high N should not reach their photosynthetic capacity while the respiration rate remains almost invariant between morning and midday conditions. In the present study, sensitivity to environmental conditions varied between the cultivars because of their contrasting stomatal behaviors (Figs. 4 and 5), which determined the gap between potential and operational photosynthetic carbon gains (Fig. 6). A high canopy N content may pose a high risk, although it is a prerequisite to increase the potential return.

‘Haying-off’ may be a similar and extreme case of the high risk associated with high canopy N. ‘Haying-off’, which has been reported in Australia, is the premature ripening of crops that leads to low grain yield and high grain protein concentration due to low post-anthesis rainfall. According to van Herwaarden *et al*. (1998), ‘haying-off’ is caused by a reduction in post-anthesis photosynthetic assimilation and is prominent in canopies growing vigorously and consuming more water, which implies that high canopy N has the potential risk of utilizing large amounts of water and exhibiting high respiration rates.

The role of photosynthesis in increasing crop yields has been debated over the years (e.g., Long *et al*. 2006). Several recent studies have emphasized the importance of operational photosynthesis rather than photosynthetic rates under suitable conditions (Murchie *et al*. 2018) and the results of the present study support the view. When photosynthetic rates of leaves were measured by introducing ambient air into the gas chamber so that the rates reflected *in situ* values, a positive relationship was observed between photosynthetic rate and grain yield in Mexican spring wheat cultivars (*N* = 8; Fischer *et al*. 1998), wheat cultivars under warm, irrigated, and dry conditions in Mexico (*N* = 16; Reynolds *et al*. 2000), and Chinese winter wheat cultivars (*N* = 18; Jiang *et al*. 2003). In studies with gas exchange in environmentally controlled gas chambers, significant correlations have also been observed between grain yield and leaf photosynthetic rates among modern UK cultivars, synthetic derivatives, landraces [*N* = 15; Gaju *et al*. (2016)], and Indian elite cultivars (*N* = 30; Nehe *et al*. 2020). However, Driever *et al*. (2014) reported that a high photosynthetic rate was not correlated with grain yield based on the analysis of a large genotypic population in winter wheat (*N* = 64). When field conditions during the growth period were suitable and operational photosynthesis was close to the values measured under controlled conditions, there should be a strong relationship between the photosynthetic rate and grain yield. When not suitable, the major determinant of grain yield should be operational photosynthesis under stressed conditions and respiratory losses, leading to the absence of correlation. Because wheat yield in the UK was the lowest over the 1990–2020 period when Driever *et al*. (2014) experimented in 2011/12 (FAO 2022), field conditions should not be suitable for the leaves to achieve their potential photosynthetic capacity. Shimoda and Sugikawa (2020) demonstrated that exposure of YC and KH cultivars to solar radiation immediately after anthesis had a considerable impact on grain yield, suggesting the importance of post-anthesis photosynthetic carbon gain on yield.

### Distinct photosynthetic strategies of elite cultivars and their implication for breeding to mitigate against climate change

Intensive gas exchange measurements have revealed that elite wheat cultivars exhibit contrasting photosynthetic performance, characterized mainly by their stomatal behavior. The present findings highlighted that assessment of photosynthesis under conditions almost similar to those in the field is necessary because photosynthetic performance may differ from that expected from leaf N content and/or that evaluated under controlled environmental conditions. Field phenotyping of crops based on the functional aspects of plants, particularly photosynthesis and transpiration, as well as their structural traits, should be promoted. For example, previous studies indicated a negative correlation between remotely sensed canopy temperature and wheat yield (e.g., Idso *et al*. 1977; Fischer *et al*. 1998). In the present study, the high-yielding cultivar exhibited a higher stomatal conductance than the low-yielding cultivar, implying a low canopy temperature due to the cooling effect of transpiration. In addition, preliminary data suggested that leaf temperature tended to be lower in KH than in YC (Fig. S8). The results indicate that temperature-based phenotyping may be effective for selecting high-yielding lines. However, the lines selected by this method could be vulnerable to drought stress due to their high water requirements. The breeding method may also lead to a decrease in the genetic diversity of the population and loss of traits that are useful for enhancing water use efficiency.

In the context of climate change, loose stomatal regulation, such as that in the KH cultivar could have both negative and positive effects. A potential positive effect is the lowering of the leaf temperature through transpirational cooling. When the leaf temperature exceeds an optimum temperature range for photosynthesis, the cooling effect promotes leaf photosynthesis. Although Nagai & Makino (2009) observed that the net photosynthetic rate measured at the growth temperature remained almost constant in wheat and rice within a wide range of the growth temperature, it is not clear whether this compensation will be effective under the projected temperature increase of up to 4–6 K, depending on the region and socioeconomic pathway (Masson-Delmotte *et al*. 2021). A plausible negative effect is the mitigation of drought tolerance. Establishing a drought-resilient wheat production system is a major concern in most wheat-producing regions. Abundant fresh water due to intensive snowfall and summer precipitation received at our study site appeared to emphasize the positive aspect of the loose stomatal regulation in KH and caused a higher grain yield. The effects of drought under the present climate conditions could be negligible because this cultivar has not previously exhibited a notable reduction in grain yield during a dry year (Shimoda & Sugikawa 2020), and the yield tended to be lower in wet years than in dry years (Murakami *et al*. 2021). However, climate models suggest a change in the pattern of rainfall and an increase in the chances of intense drought events (Masson-Delmotte *et al*. 2021). Excessive water loss through transpiration could surpass the amount of available soil water during the dry periods under future climate conditions. Models of leaf thermodynamics and hydrological conditions are required for the quantitative assessment of climate risks.

The total amount of grain N is mainly determined by N stored in vegetative organs at anthesis; therefore, increased photosynthetic carbon gain in high-N canopies is required to simultaneously achieve high grain yield and protein concentration. The difference between the potential and operational photosynthetic rates of leaves at a given N level is a major issue that needs to be addressed. In the present study, we observed that the contrasting stomatal behavior between the two cultivars was a major determinant of operational photosynthesis (Figs. 6 and 7). Previous studies have identified key proteins involved in the regulation of the stomatal opening and closing (Shimazaki *et al*. 2007). Incorporating related genes may be effective in improving canopy photosynthetic carbon gain in the low-yielding cultivar. In addition to genetic approaches, precision fertilization control in combination with weather forecasts is required to obtain high returns while minimizing risk. If meteorological conditions during the grain-filling period are suitable for leaves to achieve their potential photosynthetic capacity, additional N application may be an effective practice. Several studies have revealed that post-anthesis N uptake determines grain protein deviation (grain yield × grain protein concentration) (Monaghan *et al*. 2001; Bogard *et al*. 2010), and late N application may be effective if the weather permits. If the conditions are not suitable, high and/or late N application may not be effective because of an increase in canopy respiration. Incorporating numeric weather forecasts into agricultural management should be promoted to support decision-making.

## Acknowledgments

We are grateful to Shigeko Komine, Takahiro Mikuni, and Makoto Yamazaki for their careful plant management. We thank Atsushi Yagioka and Kenji Kimiwada for their technical support in the CN analysis. This study was financially supported by the Japan Society for the Promotion of Science (KAKENHI 19H00963, 20K15630, and 20H03112).

