## Supplemental Figures for "Trade-off between grain yield and protein concentration is modulated by canopy photosynthesis in Japanese wheat cultivars"

### Supporting information

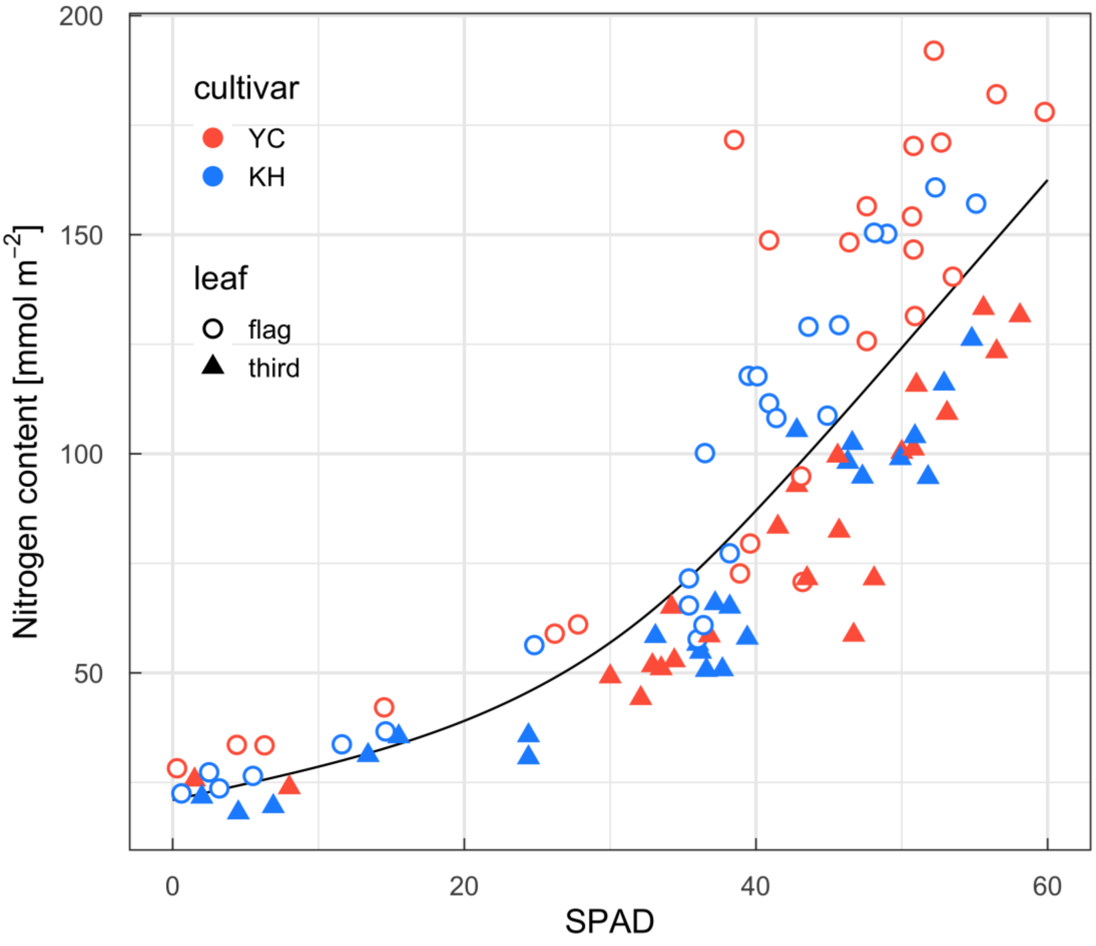


Fig. S1. A relationship between leaf nitrogen content and SPAD values. A spline curve was calculated by pooling data for the two cultivars and two leaf levels.

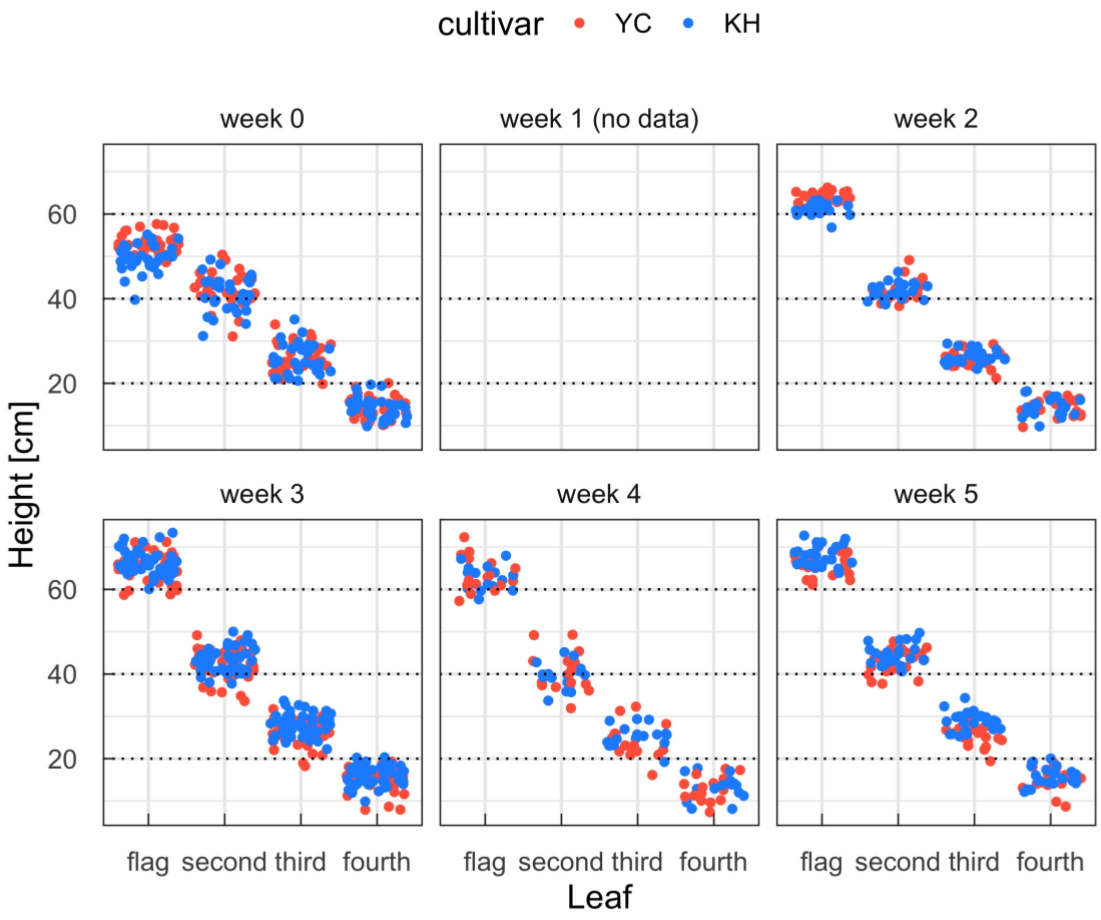


Fig. S2. Heights from the ground to the leaf bases at different growth stages.

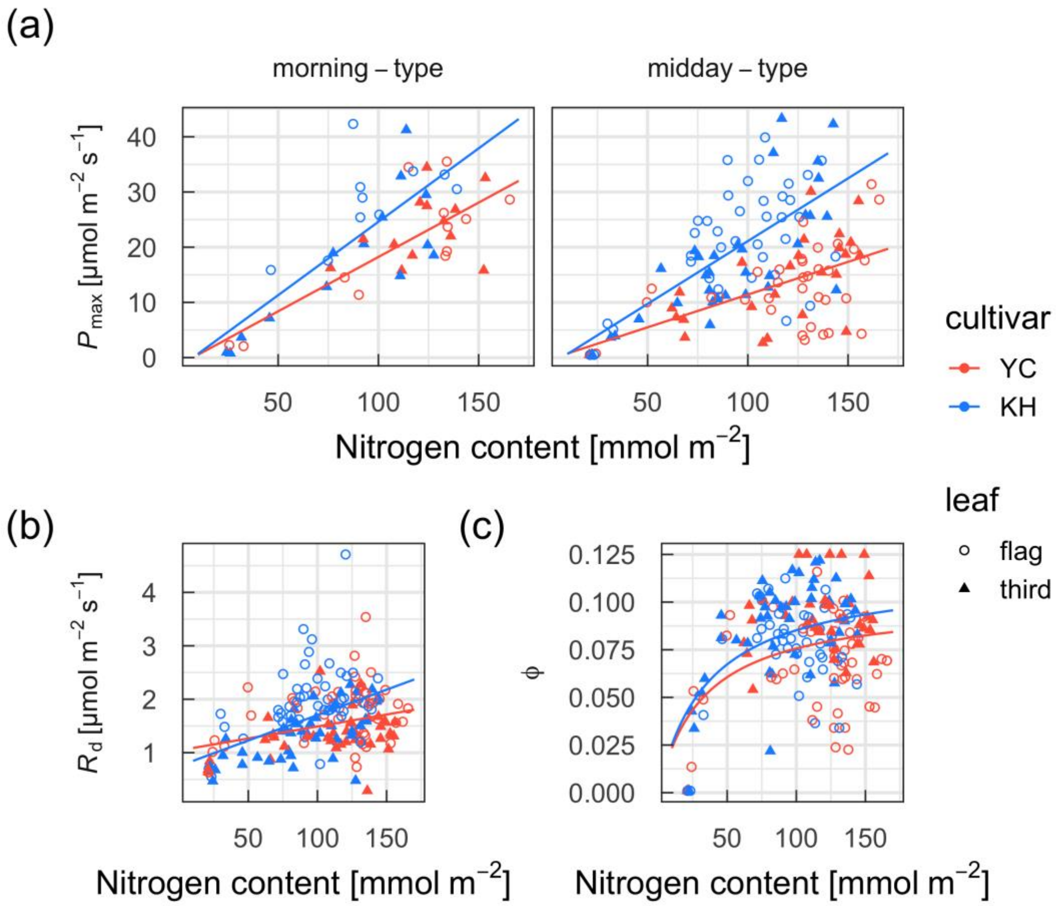


Fig. S3. Relationships among maximum photosynthetic rate ($P_{max}$), dark respiration rate ($R_d$), and the initial slope of the light response curve (ϕ) and leaf nitrogen content of ‘Yumechikara’ (YC) and ‘Kitahonami’ (KH) wheat cultivars. Morning- and midday-type regression lines for $P_{max}$ were determined separately.

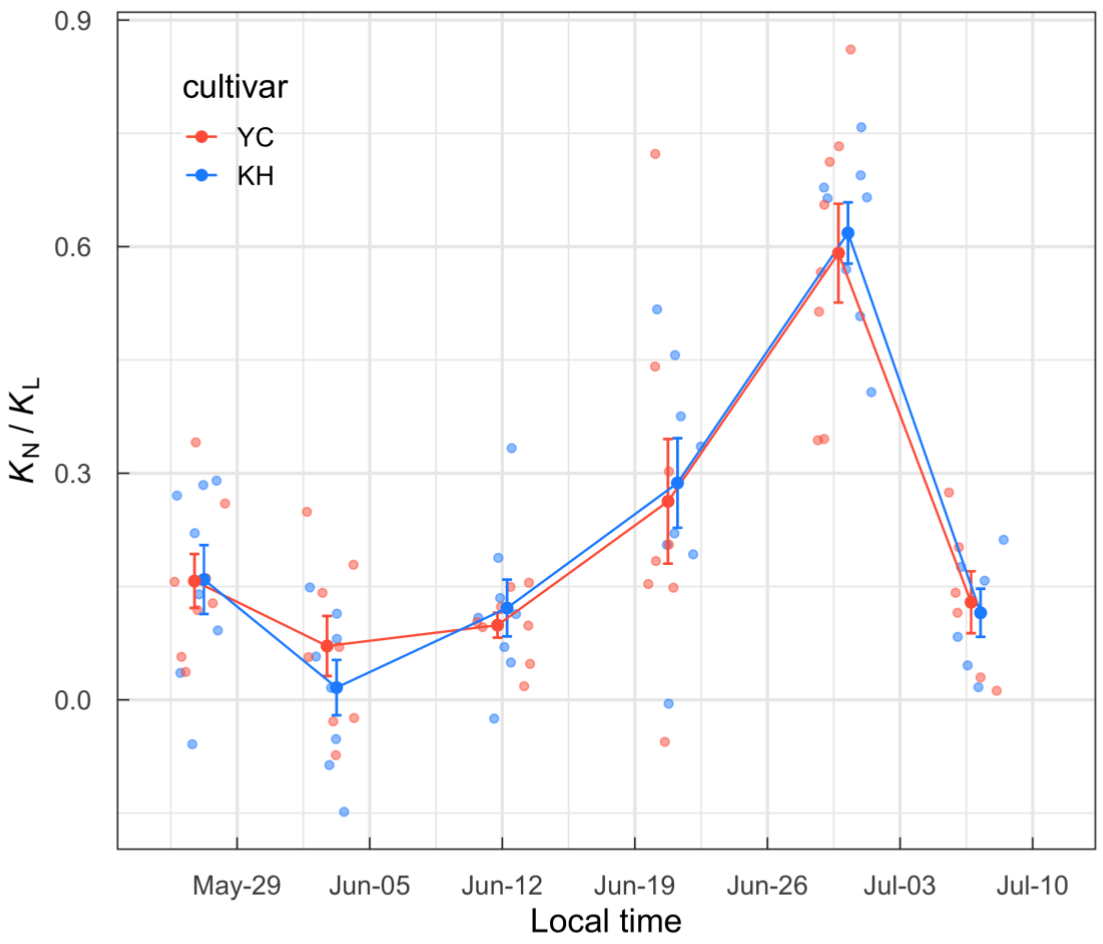


Fig. S4. Ratios of the nitrogen extinction coefficient ($K_N$) to the light extinction coefficient ($K_L$) of ‘Yumechikara’ (YC) and ‘Kitahonami’ (KH) wheat cultivars as a function of date. Values of eight replicates (small circles), their means (large circles) and standard errors (error bars) are shown."

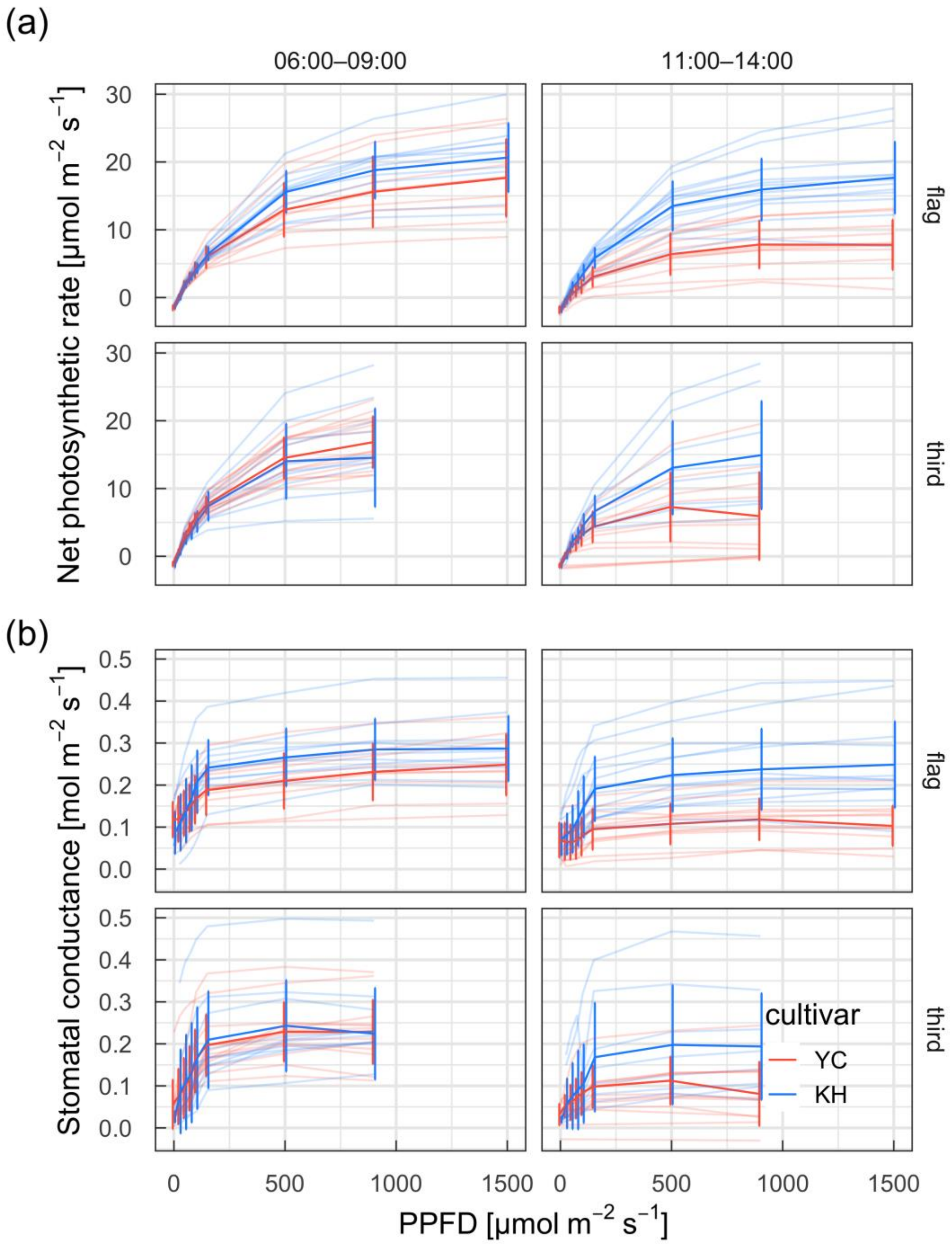


Fig. S5. Net photosynthetic rates and stomatal conductance of flag and third leaves of ‘Yumechikara’ (YC) and ‘Kitahonami’ (KH) wheat cultivars measured in the morning (06:00–09:00) and at midday (11:00–14:00) as a function of photosynthetic photon flux density (PPFD). Data obtained in the last week (week 6) were not used because of progressive leaf senescence. Values of samples (thin lines), their means (thick lines), and standard deviations (error bars) are shown ($N$ = 6–14).

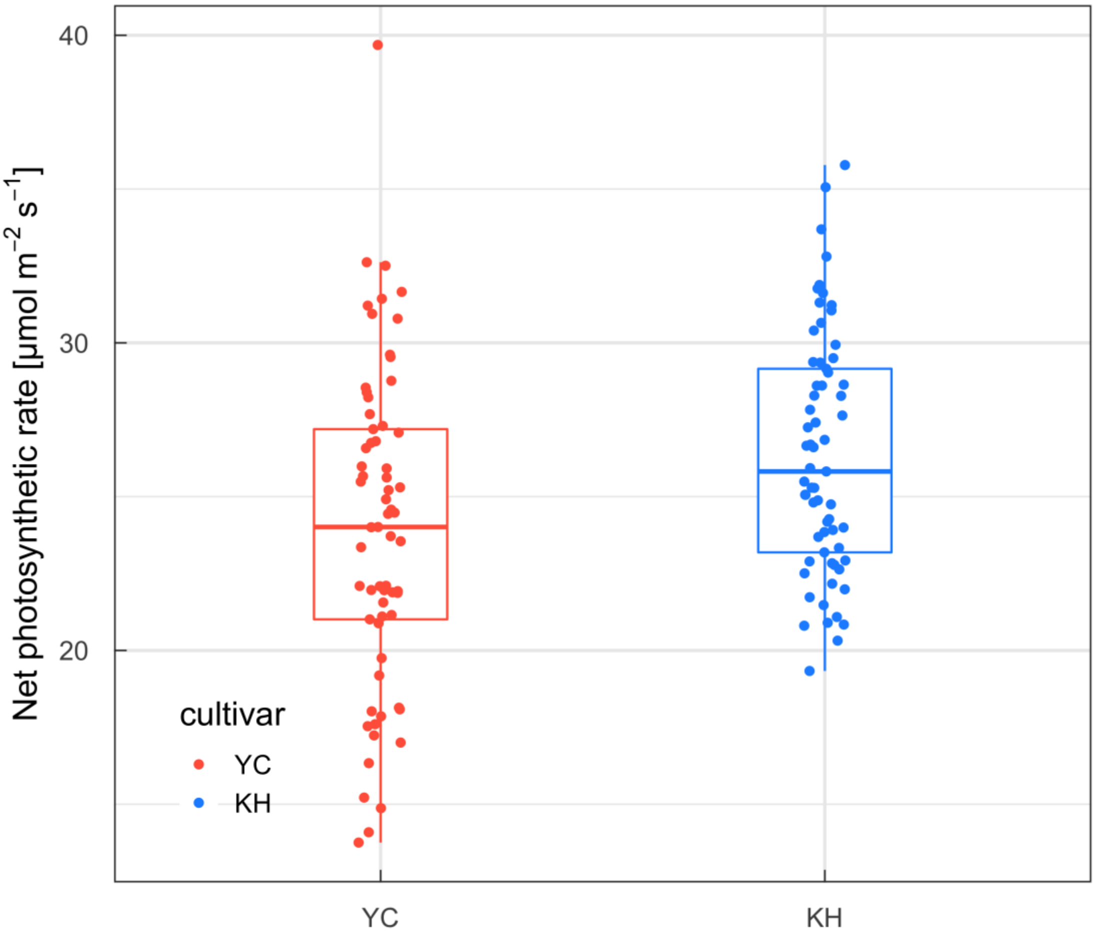


Fig. S6. Net photosynthetic rates of flag leaves of ‘Yumechikara’ (YC) and ‘Kitahonami’ (KH) wheat cultivars measured under *in situ* conditions on 2022-06-14. Measurements were carried out using a closed gas exchange system (MIC-100-S1; Masa International Corp., Kyoto, Japan) by introducing ambient air and the leaves irradiated by incident sunlight.

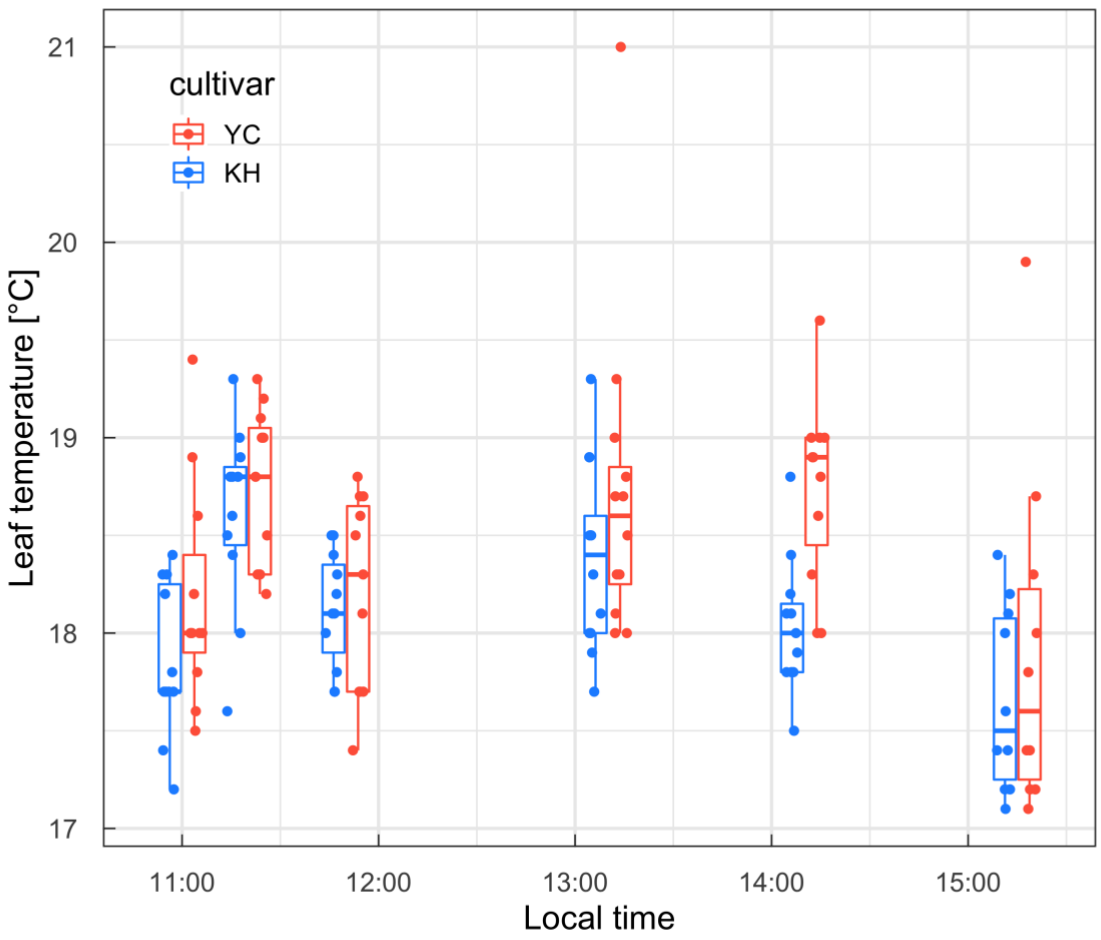


Fig. S7. Leaf temperature of flag leaves of ‘Yumechikara’ (YC) and ‘Kitahonami’ (KH) wheat cultivars measured under *in situ* conditions on 2022-06-14. Measurements were carried out using an infrared thermometer (IT-545; Horiba Co., Ltd., Kyoto, Japan).
